# The BASP1 transcriptional repressor modifies chromatin through lipid-dependent and lipid-independent mechanisms

**DOI:** 10.1101/2022.02.15.480538

**Authors:** Alexander J. Moorhouse, Amy E. Loats, Kathryn F Medler, Stefan G. E. Roberts

## Abstract

The transcriptional corepressor BASP1 requires N-terminal myristoylation for its activity and functions through interactions with nuclear lipids. Here we determine the role of BASP1 lipidation in histone modification and the modulation of chromatin accessibility. We find that the removal of the active histone modifications H3K9ac and H3K4me3 by BASP1 requires the N-terminal myristoylation of BASP1. In contrast, the placement of the repressive histone modification, H3K27me3, by BASP1 does require BASP1 lipidation. RNA-seq and ATAC-seq analysis finds that BASP1 regulates the activity of multiple transcription factors and induces extensive changes in chromatin accessibility. We find that ∼50% of BASP1 target genes show lipidation-dependent chromatin compaction and transcriptional repression. Our results suggest that BASP1 elicits both lipid-dependent and lipid-independent functions in histone modification and transcriptional repression. In accordance with this, we find that the tumor suppressor activity of BASP1 is also partially dependent on its myristoylation.

## Introduction

BASP1 was initially identified as a cytoplasmic signaling protein in neuronal cells (Mosevitsky, 2005) but has since been found to be widely expressed and localizes to the nucleus in several cell types via a canonical nuclear localization sequence (Carpenter et al., 2004; Goodfellow et al., 2011; Toska and Roberts, 2014). A role for BASP1 in transcriptional regulation was first identified as a cofactor for the Wilms’ tumor 1 protein WT1 (Carpenter et al., 2004). BASP1 binds to WT1 and converts it from a transcriptional activator to a repressor. BASP1 has subsequently been found to act as a repressive cofactor for other transcriptional regulators including MYC (Hartl et al., 2009), ER*α* (Marsh et al., 2017) and YY1 (Santiago et al., 2021) suggesting that it is broadly deployed in transcription control (Hartl and Schneider, 2019). Our understanding of how BASP1 regulates transcription is limited and has, to date, largely been studied as a transcriptional corepressor of WT1. BASP1 is N-terminally myristoylated and we demonstrated that this lipid motif is required for its function as a transcriptional repressor (Toska et al., 2012, 2014). BASP1 also contains a consensus cholesterol binding motif adjacent to the myristoyl moiety (Epand, 2008). Indeed, both phosphatidylinositol 4,5-bisphosphate (PIP2) and cholesterol are recruited to promoter regions and are required for transcriptional repression by BASP1 (Toska et al., 2012, 2014; Loats et al., 2021). The myristoylation of BASP1 is required for interaction with both PIP2 and cholesterol (Toska et al., 2012; Loats et al., 2021). PIP2 assists in the recruitment of HDAC1 to mediate deacetylation of histone H3K9 and direct transcriptional repression (Toska et al., 2012).

Several studies have demonstrated that BASP1 acts as a tumor suppressor, for example in hepatocellular carcinoma (Tsunedomi et al., 2010), gastric cancer (Li et al., 2020) and breast cancer (Marsh et al., 2017). Consistent with its tumor suppressor activity, BASP1 has a role in maintaining the differentiated state and is required to maintain the functional differentiated state of taste receptor cells in mice (Gao et al., 2019). BASP1 can also block the transformation of fibroblasts by v-myc (Hartl et al., 2009). Moreover, BASP1 interferes with the action of the Yamanaka proteins in the induction of pluripotent stem cells by repressing WT1 target genes (Blanchard et al., 2017). Taken together, current studies suggest that BASP1 plays a major role in maintenance of the differentiated state through the modulation of the function of several transcriptional activator proteins.

The mechanism of action of BASP1 as a transcription factor is poorly understood, but its physiological activities suggest extensive roles in gene regulation. In this study we set out to determine how BASP1 regulates transcription through histone modifications and the roles of the myristoyl motif of BASP1. We find that lipidation of BASP1 plays a selective role in its action as a transcription factor and is required for specific histone modification leading to modulation of the chromatin environment in order to elicit specific transcriptional effects. Our results suggest that BASP1 plays a widespread role in regulating the chromatin environment and transcription control.

## Results

### BASP1 mediates the removal of active histone modifications and the placement of repressive H3K27me3

K562 cells express endogenous WT1 but do not express BASP1 (Goodfellow et al., 2011). As we have shown before, the introduction of BASP1 into K562 cells (B- K562 cells) leads to the transcriptional repression of the WT1 target genes AREG and VDR compared to the control K562 cells (V-K562 cells; Figure 1A; Goodfellow et al., 2011; Toska et al., 2012, 2014; Loats et al., 2021). We next performed chromatin immunoprecipitation (ChIP) to detect WT1, BASP1, and a selection of specific histone modifications at the promoter regions of the WT1 target genes AREG and VDR (Figure 1B). WT1 was present at the promoter region of the AREG and VDR genes in both control V-K562 cells and B-K562 cells, while BASP1 was present at the promoter region only in B-K562 cells. Analysis of H3K9ac and H3K4me3, indicators of transcriptionally active promoter regions, demonstrated that these histone modifications are removed when BASP1 is recruited to the promoter in B- K562 cells and the genes are transcriptionally repressed. These data are consistent with our previous studies of BASP1 function in histone modification (Toska et al., 2012; Loats et al., 2021). We also analyzed the H3K27me3 histone modification associated with transcriptional repression. We found that BASP1 induces the trimethylation of H3K27 at both the AREG and VDR promoter regions. Taken together, these results demonstrate that BASP1 functions by both the removal active histone modifications (H3K9ac and H3K4me3) and the placement of the repressive histone mark H3K27me3.

**Figure 1.**
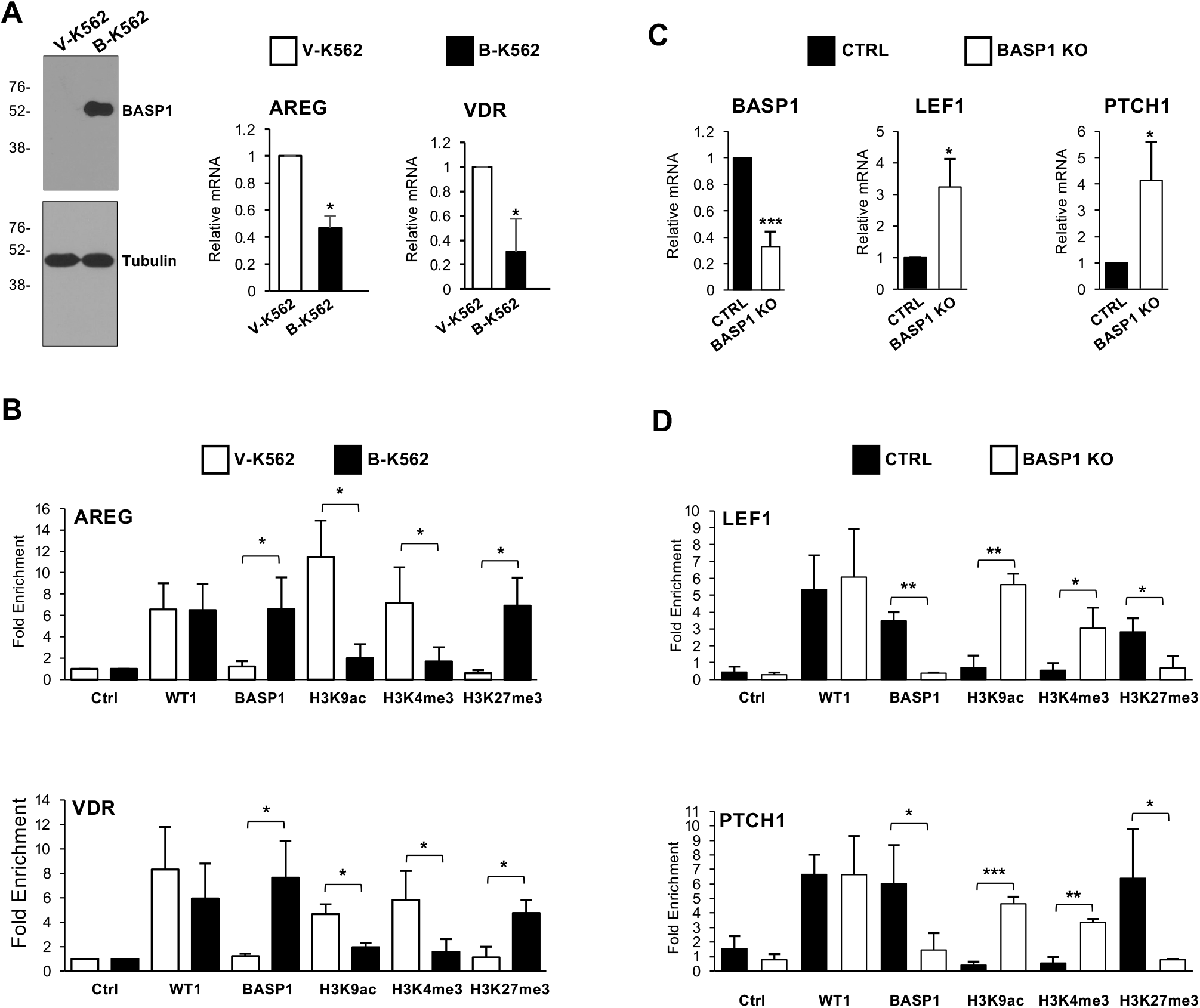
BASP1 directs removal of active histone modifications H3K9ac and H3K4me3 and the placement of repressive H3K27 trimethylation. (A) Immunoblotting of extracts prepared from V-K562 and B-K562 cells to confirm BASP1 expression. *β*-tubulin immunoblotting was performed as a loading control. Molecular weight markers are shown at left (kDa). cDNA was prepared from V-K562 cells and B-K562 cells and expression of AREG and VDR quantitated relative to GAPDH. Data are Standard Deviation from the mean (SDM) for three independent experiments. *=p<0.05 by students t-test. (B) V-K562 or B-K562 cells were subjected to chromatin immunoprecipitation (ChIP) with control (ctrl) antibodies or antibodies against WT1, BASP1 or the histone modifications indicated. Data are presented as fold-enrichment over a control genomic region and error bars are SDM of three independent experiments. *=p<0.05 by students t-test. (C) Krt8-BASP1-CRE mice were treated with tamoxifen for 8 days and then 7 days later taste buds were isolated and RNA prepared. cDNA was then use to monitor expression of BASP1, LEF1 and PTCH1 compared to GAPDH. Error bars are SDM for three independent experiments. *=p<0.05 by students t-test. (D) Mice were treated as in part C but isolated taste cells were subjected to chromatin immunoprecipitation with the antibodies indicated. Error bars are SDM for three independent experiments. *=p<0.05, **=p<0.01 and ***=p<0.005 by students t-test.

We have previously used a conditional BASP1 mouse to demonstrate that BASP1 transcriptionally represses the WT1 target genes LEF1 and PTCH1 in taste receptor cells (Gao et al., 2019). A floxed BASP1 mouse was crossed with a Krt8-Cre-ER mouse to delete BASP1 in the fully differentiated Krt8-expressing taste cells. We used this model to perform ChIP to analyse BASP1-dependent histone modifications at the LEF1 and PTCH1 promoters in taste receptor cells. Consistent with our previous studies, knockout of BASP1 expression in taste cells leads to the upregulation of LEF1 and PTCH1 expression (Figure 1C). ChIP analysis confirmed that WT1 was present at the promoter region of the LEF1 and PTCH1 genes in both the control and BASP1 KO taste cells while BASP1 was only present at the promoter regions in the taste receptor cells from control mice (Figure 1D). Analysis of H3K9ac, H3K4me3 and H3K27me3 demonstrated that loss of BASP1 leads to the accumulation of the active histone modifications H3K9ac and H3K4me3, and the removal of the H3K27me3 repressive modification. These findings, using an *in-vivo* model of BASP1 function, confirm and extend our analysis in K562 cells demonstrating that BASP1 directs the removal of the active histone modifications H3K9ac and H3K4me3, and the placement of repressive H3K27me3.

### Myristoylated BASP1 is recruited to gene promoters

Our previous work demonstrated that the mutation BASP1-G2A, which prevents N-terminal myristoylation of BASP1, leads to loss of transcriptional repression function (Toska et al., 2012, 2014). Although total cellular BASP1 is stoichiometrically N-terminally myristoylated (Mosevitsky, 2005), it has not yet been demonstrated that this lipidated form of BASP1 is present within the nucleus and is directly recruited to gene promoter regions. We therefore used a click chemistry approach to analyze the myristoylation status of nuclear BASP1 in K562 cell line derivatives. V-K562 and B- K562 cells along with G-K562 cells (expressing BASP1-G2A) were incubated in lipid-free media supplemented with myristic acid alkyne that is utilized by cells as a substrate for N-terminal myristoylation (Wright et al., 2015). Nuclei were then prepared as we described before using a method that avoids contamination with either endoplasmic reticulum or golgi complex (Loats et al., 2021). Using click chemistry, the alkyne-myristoyl moiety was crosslinked to azide-PEG3-biotin which was then detected using a streptavidin-linked fluorophore. BASP1 was simultaneously probed using immunocytochemistry. In control V-K562 cells, myristoyl was detected within the purified nuclei of K562 cells and was particularly evident at the nuclear membrane (Figure 2A). This staining pattern is consistent with the known N-terminal myristoylation of lamins (Linde and Stick, 2010). In B-K562 cells there was similar staining at the nuclear periphery and also an enhanced intranuclear staining. BASP1-G2A showed reduced colocalisation with myristoyl compared to wtBASP1 (Pearson quantitation of the proportion of BASP1 signal that overlaps with myristoyl is shown in Figure 2B).

**Figure 2.**
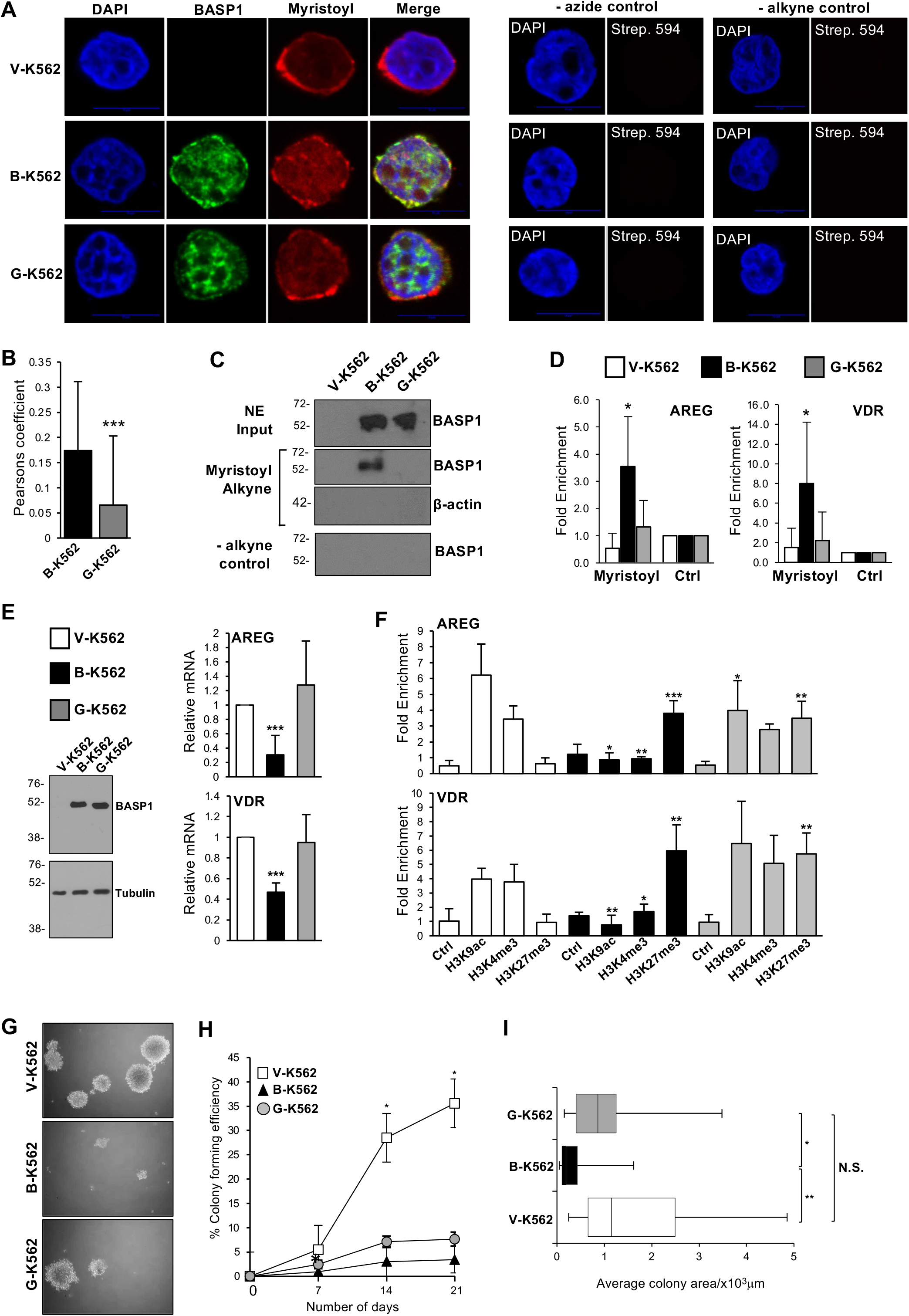
The role of myristoylation of BASP1 in transcription, histone modification and tumorigenesis. (A) The indicated cell lines were incubated in lipid-free media in the presence of 10μg/ml myristic acid alkyne for 20 hours. Nuclei were prepared and incubated with azide PEG3-biotin, then the click reaction initiated using 2mM CuBF4. Immunohistochemistry was then performed with streptavidin-linked antibodies (Myristoyl) and BASP1 antibodies. Scale bar is 10μm. Control assays were performed that lacked azide PEG3-biotin (-azide control) or myristic acid alkyne (- alykyne control) and are shown at right. (B) Quantification of the colocalisation of BASP1 and Myristoyl in B-K562 and G-K562 nuclei was analyzed using the Pearson’s correlation coefficient. ***=p<0.005 following students t-test comparing Pearson’s values from BK and GK nuclei over 3 independent experiments (n=67). (C) V-K562, B-K562 and G-K562 cells incubated with ethanol (-alkyne control) or 10μg/ml alkyne-Myristic acid as above and subjected to a click chemistry reaction. Nuclear extracts were prepared and precipitation was performed with streptavidin beads. Immunoblotting was then performed with either BASP1 antibodies or control β-actin antibodies. Molecular weight marker in kDa shown to left of each blot. (D) The cell line derivatives were treated in part A and following the click chemistry reaction, ChIP was performed with either streptavidin-linked beads (Myristoyl) or control beads (Ctrl). Data are shown as fold enrichment of Myristoyl at the AREG and VDR promoters in V-K562, B-K562 and G-K562 cells compared to the alkyne-free control precipitation. Error bars SDM with *=p<0.05 by student’s t test comparing B-K562 or G-K562 with control cell line V-K562. (E) Expression of the WT1 target genes AREG and VDR was monitored relative to GAPDH in V-K562, B-K562 and G-K562 cells. Immunoblotting was performed to confirm the expression of the BASP1 derivatives. (F) V-K562, B-K562 and G-K562 cells were subjected to ChIP analysis with antibodies against H3K9ac, H3K4me3, H3K27me3 or control. Data are presented as fold enrichment at the AREG and VDR promoters compared to a control genomic region. (G) Representative images of V-K562, B-K562 and G-K562 cells seeded into agar dishes for 21 days. (H) Colony formation efficiency was assessed at days 7,14 and 21 for V-K562, B-K562 and G-K562 cells. Error bars are SDM of 3 independent experiments. (I) Box and Whisker plot of average colony area after 21 days growth of V-K562, B-K562 and G-K562 cells in agar. *=p<0.05, **=p<0.005, n.s. (no significant difference) by student’s t-test comparing 3 independent experiments.

We next used click chemistry combined with immunoprecipitation to determine the association between BASP1 and nuclear myristoyl. V-K562, B-K562 and G-K562 cells were cultured in the presence of myristic acid alkyne as above for 48 hours and, following the click chemistry reaction with azide-PEG3-biotin, nuclear protein extracts were prepared then immunoprecipitation was performed with streptavidin-coated beads. wtBASP1, but not BASP1-G2A, co-immunoprecipitated with the alkyne- myristoyl (Figure 2C). We have previously combined click chemistry with ChIP (Click- ChIP; Loats et al., 2021) and therefore used this technique to determine if myristoyl is recruited to the promoter region of WT1 target genes. We found that myristoyl associates with the AREG and VDR promoters at a significantly increased level when wtBASP1 is also present at the promoter (Figure 2D; compare V-K562 with B-K562), but at a reduced level in K562 cells that express BASP1-G2A (G-K562). These data demonstrate that detection of myristoyl at the promoter region of WT1 target genes is dependent on BASP1 and that this association requires an intact G2 target site of myristoylation in BASP1.

### Myristoylation of BASP1 is required for the removal of active chromatin modifications but not for the placement of repressive H3K27me3

Our data so far have demonstrated that myristoylated BASP1 is present at the promoter region of WT1 target genes. Our previous work (Toska et al., 2012, 2014), confirmed in Figure 2E, has shown that BASP1-G2A lacks transcriptional corepressor activity. We demonstrated before that BASP1-G2A is defective in deacetylation of H3K9 (Toska et al., 2012). We therefore tested if BASP1-G2A was able to mediate demethylation of H3K4me3 or the placement of repressive H3K27me3. ChIP analysis confirmed that BASP1-G2A was defective in the removal of the active H3K9ac mark (Figure 2F). We also found that BASP1-G2A was defective in the removal of H3K4me3. Thus, myristoylation of BASP1 is required for the removal of active histone modifications. In contrast, BASP1-G2A was still able to support the placement of repressive H3K27me3 at both the AREG and VDR promoters. Thus, the N-terminal myristoylation of BASP1 is required to mediate removal of the active histone modifications, but is not required to place the repressive H3K27me3 modification. These findings suggest a dual mechanism of lipidation-dependent and lipidation-independent transcriptional repression by BASP1.

### Myristoylation of BASP1 plays a role in the tumor suppressor activity of BASP1

BASP1 acts as tumor suppressor in several cell types and slows the growth of K562 cells (Goodfellow et al., 2011; Toska et al., 2014). We therefore tested the effects of wtBASP1 and BASP1-G2A on the formation of anchorage independent colonies of K562 cells in soft agar assays (Figure 2G). The number of V-K562, B- K562 and G-K562 colonies greater than 50 cells in size were counted over a three-week period (Figure 2H). Expression of wtBASP1 led to a significant decrease in the colony forming ability of K562 cells. Measurement of the average area of colonies revealed that, as well as forming fewer colonies, B-K562 cells form significantly smaller colonies (Figure 2I). Expression of BASP1-G2A also led to a significant decrease in the colony forming ability of K562 cells (Figure 2H). However, the average area of G-K562 colonies did not significantly differ to that of V-K562 colonies (Figure 2I). Thus, myristoylation of BASP1 is not required to suppress the formation of anchorage independent colonies but contributes to inhibiting the growth of the established colonies. We conclude that the myristoylation of BASP1 plays a partial role in its tumor suppressor function in K562 cells.

### Genome-wide analysis of the requirement for BASP1 myristoylation in transcriptional repression

Our results so far demonstrate that BASP1 directs histone modifications that are both dependent (H3K9 deacetylation and H3K4 demethylation) and independent (H3K27 methylation) on the N-terminal myristoylation of BASP1. In addition, myristoylation of BASP1 is partially required for its tumor suppressor activity. Taken together, these findings suggest that non-myristoylated BASP1 is likely to retain partial activity in transcriptional regulation that we have not observed here or before (Toska et al., 2012, 2014) in targeted gene analysis. We therefore determined the gene expression profiles of B-K562, G-K562 and control V-K562 cells using RNA-seq. Comparing the data from V-K562 and B-K562 cells (Figure 3A) supports our previous genome-wide analysis with 3064 genes regulated by BASP1 (2165 genes repressed and 899 genes activated; Goodfellow et al., 2011). As before, the majority of the known WT1 target genes that show altered expression were transcriptionally repressed (70.3% of the 105 WT1 target genes that changed significantly (padj 0.05); Goodfellow et al., 2011; Figures 3B and 3C). Our data further show that BASP1-G2A is partially defective in transcriptional repression when compared to wtBASP1, repressing 58.8% of the genes that are repressed by wtBASP1 (Figure 3A). This was reduced to 45.7% when considering only WT1 target genes (Figures 3B and 3D). In fact, BASP1-G2A activated many of the target genes that are repressed by wtBASP1 (Figure 3D).

**Figure 3:**
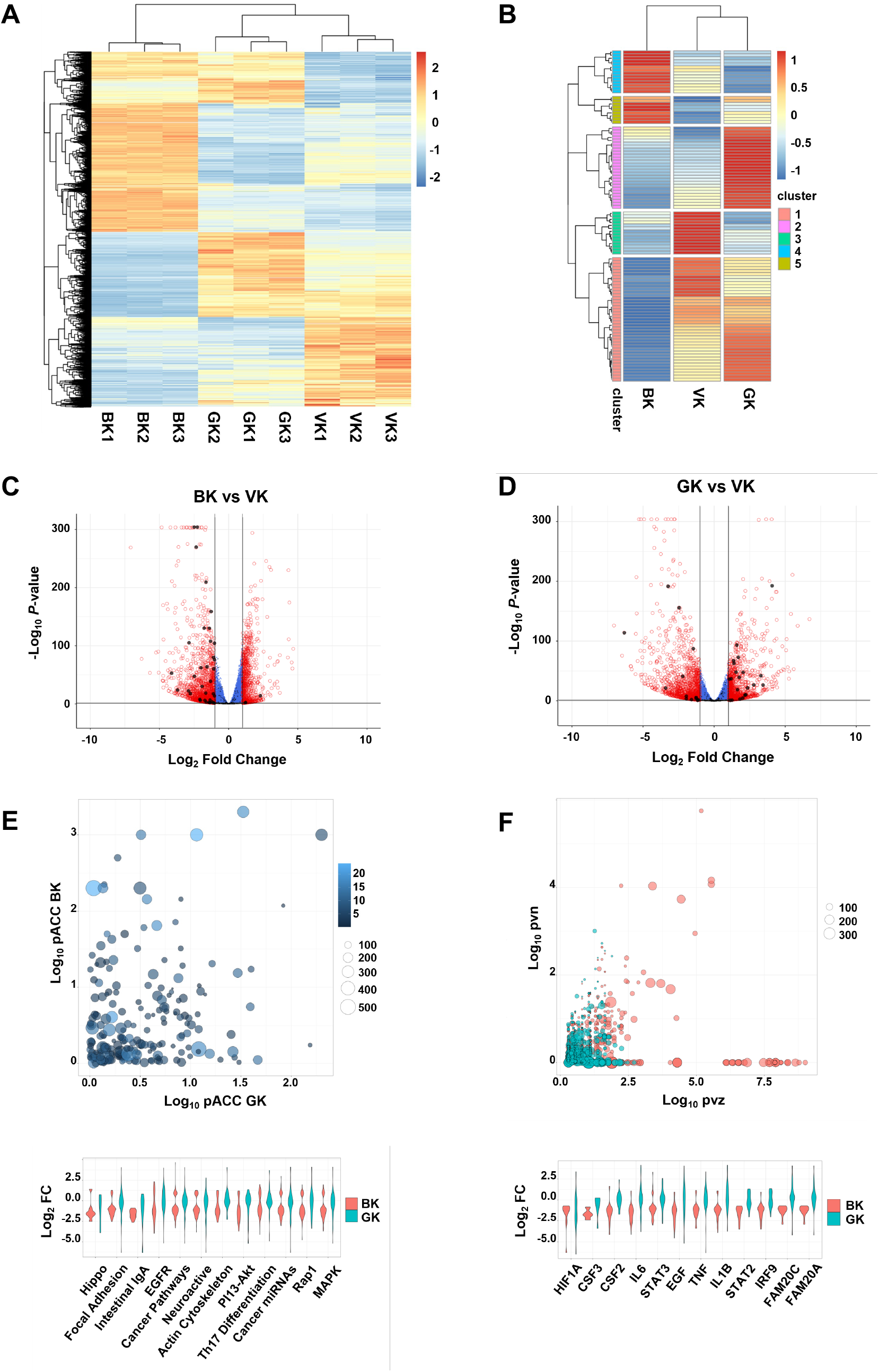
BASP1 elicits lipidation-dependent and lipidation-independent transcriptional regulation. (A) Gene expression is shown for genes at padj ≤ 0.05 and fold-change ≥ 1 for BK vs VK for all genes. Data from three independent RNAseq experiments for V-K562, B- K562 and G-K562 cells is shown. (B) As in part A but only analysis of known WT1 target genes is shown. Signals are the average of three independent RNAseq experiments. (C) Differential gene expression is shown in volcano plots for B-K562 cells vs V-K562 cells. WT1 target genes are shown black. (D) As in part C but comparing V-K562 cells with G-K562 cells. (E) Differentially regulated gene networks are plotted for BK and GK, compared with VK: Canonical pathways are positioned by -Log10 perturbation accumulation (pAcc) for B-K562 cells (y axis) and for G-K562 cells (x axis). Pathways are sized by the number genes in each pathway and coloured by pathway over-representation (pORA) for B-K562 cells vs V-K562 cells. (F) Upstream regulators inhibited in B-K-562 cells are positioned by -Log10 P values for the number of targets consistent with inhibition (pvn) and regulator z-score (pvz), balloons are sized by the total number of target genes for each regulator. The following thresholds were applied to E: -Log10 pAcc ≥ 2, -Log10 pvn ≥ 1.5, and to F: -Log10 pvz ≥ 2.5, violin plots are ordered left to right by -Log10 pAcc and, -Log10 pvz, respectively and show fold-change values for those networks in B-K562 cells and G-K562 cellks, vs V-K562 cells.

Pathway enrichment analysis of genes repressed by wtBASP1 reveal several networks (canonical pathways and upstream regulators) that are enriched for significantly repressed differentially expressed genes in B-K562 cells, and further demonstrate their derepression in G-K562 cells (Figures 3E and 3F). We also identified other networks downregulated in B-K562, but not G-K562 cells, including those involved in the response to the H3K27 Demethylase Inhibitor GSK-J4, the selective estrogen receptor modulator Tamoxifen, the extra-cellular space and pre-ribosome cellular processes (Figure S1A). Together these data show the repressive effects of wtBASP1 are partially lost with the BASP1-G2A mutant derivative, specifically, the BASP1-dependent downregulation of transcription factors, upstream regulators, drug responses and canonical pathways genome wide (Figures 3E, 3F, S1A). BASP1 was also found to regulate the function of a number of transcription factors, in particular MYC, RELA, TFAP2A/C, YY1 and CTCF (Figure S1B). In the case of MYC, the effect of wtBASP1 was both activation and repression and a similar profile for MYC target genes was observed with K562 cells expressing BASP1-G2A. In contrast, wtBASP1 was predominantly repressive at RELA, TFAP2A, YY1 and CTCF target genes but BASP1-G2A elicited transcriptional activation at most genes that changed expression. Taken together the data in Figure 3 demonstrate that myristoylation of BASP1 is required for its transcriptional repressor function at ∼50% of target genes. Furthermore, BASP1 regulates the expression of the target genes of both previously described BASP1-associated transcription factors (MYC, YY1, CTCF) and newly identified targets.

### The effects of BASP1 and requirement for myristoylation on the chromatin landscape

Our targeted ChIP analysis of histone modifications (Figure 2) suggests that BASP1 likely makes extensive changes to the chromatin environment. Furthermore, the BASP1-G2A mutant derivative is defective in the removal of active chromatin modifications (H3K9ac and H3K4me3) but not the addition of repressive chromatin modification (H3K27me3). To investigate the effects of these BASP1-dependent chromatin remodelling events genome-wide, we conducted sequencing experiments to Assay for Transposase-Accessible Chromatin (ATAC-seq). Comparing control V- K562 cells with B-K562 cells the data demonstrate that chromatin accessibility is substantially altered genome-wide by wtBASP1 with 79% of the peak changes to closed chromatin compared to 21% of the changes to open chromatin (Figure 4A).

**Figure 4:**
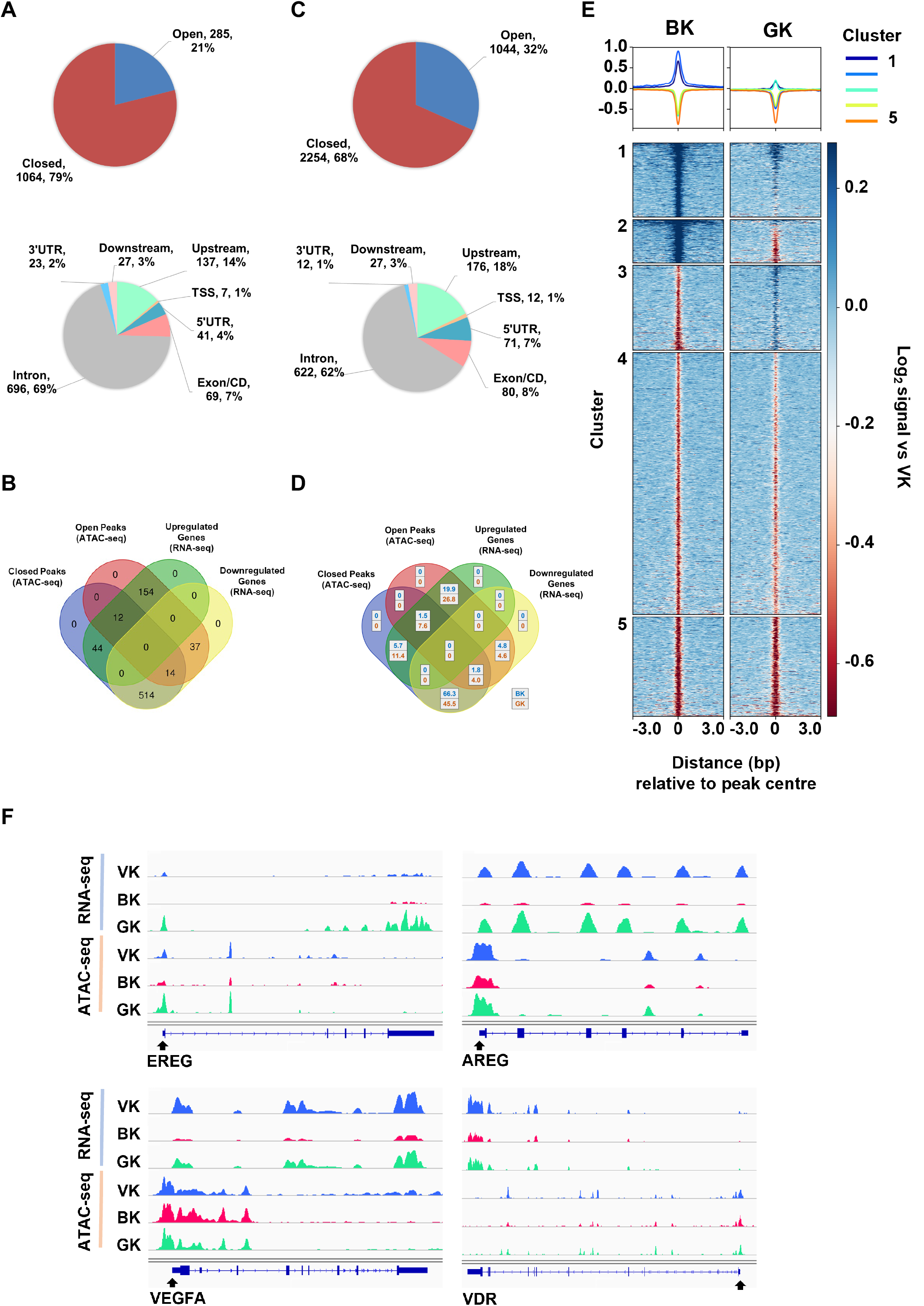
BASP1-dependent modification of chromatin accessibility is mediated by lipidation-dependent and lipidation-independent mechanisms. (A) Open and closed chromatin regions (padj ≤ 0.05 and FC ≥ 1 for B-K562 vs V- K562 cells) and their associated genomic features are shown in pie charts. (B) Chromatin regions were assigned to the closest gene within 5kb and are shown (for peaks and genes at padj ≤ 0.05 and FC ≥ 1 B-K5623 vs V-K562) as a venn diagram. (C) As in part A except that V-K562 and G-K562 cells are compared. (D) The proportion of open and closed chromatin regions at up and down regulated genes is shown for B-K562 cells and G-K562 cells. (E) Chromatin regions at padj ≤ 0.05 and fold-change ≥ 1 for B-K562 cells vs V-K562 cells are plotted as Log2 signal vs V- K562 cells for B-K562 cells and G-K562 cells. (F) Chromatin peaks and gene expression are shown for EREG, AREG, VDR and VEGFA.

These data are consistent with broad transcriptional repression by BASP1. 15% of the peak changes were within the upstream regions and transcription start sites of genes, but there were also general BASP1-dependent changes across the gene bodies. When we compared the ATAC-seq data with the RNA-seq data, we observed a strong correlation between the BASP1-dependent formation of closed chromatin within the gene proximity and transcriptional repression of the associated gene by BASP1 (Figure 4B). Specifically, 514 genes that are transcriptionally repressed by BASP1 showed BASP1-dependent formation of closed chromatin peaks. Analysis of these genes by g:profiler highlighted several categories related to the physiological functions of BASP1 including developmental processes and cell differentiation (Figure S2A).

BASP1-G2A caused more peak changes in ATAC-Seq than wtBASP1 (Figure S2B), although the ratio of changes to open chromatin versus changes to closed chromatin was increased (Figure 4C). Direct comparison of wtBASP1 with BASP1- G2A showed that 66.3% of genes affected by wtBASP1correlated with both decreased chromatin accessibility and increased transcriptional repression, while for BASP1-G2A, this only occurred for 45.5% of the genes (Figure 4D). Indeed, BASP1- G2A was either completely (Figure 4E, cluster 3) or partially (Figure 4E cluster 4) defective in inducing closed regions of chromatin at most genomic regions modified by wtBASP1. Thus, although BASP1-G2A showed a defect in driving closed chromatin at many of the sites acted upon by wtBASP1, it did not show an overall lower activity in driving the formation of closed chromatin. This is consistent with BASP1-G2A acting at several sites that are not regulated by wtBASP1.

Figure 4F shows the comparison of RNA-seq and ATAC-seq analysis across four WT1 target genes, AREG, VDR, EREG and VEGFA. All four loci show a reduction of RNA-seq signal across the genes in B-K562 cells compared to V-K562 cells, which is coincident with a decrease in the chromatin accessibility around the transcription start sites of each gene (as measured by ATAC-Seq). In G-K562 cells, AREG, VDR and EREG are not transcriptionally repressed (in-fact, BASP1-G2A activates transcription of EREG), while VEGFA is still partially repressed by BASP1-G2A. Both EREG and AREG show that BASP1-G2A fails to induce closed ATAC peaks in the vicinity of the transcription start site which is coincident with the failure of BASP1- G2A to repress transcription of these genes. BASP1-G2A produced an ATAC profile similar to wtBASP1 at the transcription start site region of the VDR gene but instead showed a reduction in ATAC peaks within the main body of the gene. Taken together, these data suggest that BASP1 transcriptionally represses WT1 target genes through the formation of closed chromatin. Furthermore, BASP1-G2A shows gene-specific effects in function, both in the regulation of chromatin compaction and in transcriptional repression.

## Discussion

BASP1 represses transcription through a novel mechanism that requires its lipidation to recruit PIP2 and cholesterol to the promoter region of target genes. The localization of lipids, including PIP2 and cholesterol, within the nucleus as well as their association with chromatin is well-documented, but their roles in nuclear processes are poorly understood (Garcia-Gil et al., 2017; Fernandes et al., 2018; Fiume et al., 2019; Barbosa and Siniossoglou, 2020; Gapa et al. 2022). Here we have presented evidence that myristoylated BASP1 is recruited to the promoter of WT1 target genes. We find that the N-terminal myristyoylation of BASP1 controls its ability to modulate the chromatin environment and is needed for the full tumor suppressor activity of BASP1.

We have previously shown that BASP1interacts with PIP2 to recruit HDAC1 to the gene promoter, which then deacetylates histone H3K9. The enzymes responsible for the BASP1-dependent demethylation of K3K4 and the trimethylation methylation of H3K27 have yet to be identified. BASP1-G2A retains the function to trimethylate H3K27 but is not able to mediate deacetylation of H3K9 or demethylation of H3K4.

This raises the possibility that BASP1-G2A leads to the formation of bivalent histone marks. Bivalent marks are associated with poised promoter regions that can either be activated or repressed (Voigt et al., 2013). The BASP1-G2A mutant derivative still causes significant changes in chromatin accessibility suggesting that the H3K27 trimethylation function is sufficient, at least for a subset of target genes, to remodel chromatin and alter gene expression. Indeed, our RNAseq data demonstrates that BASP1-G2A retains transcriptional repressor activity at ∼46% of WT1 target genes and ∼59% overall. Several previous studies have demonstrated the tumor suppressor activity of BASP1 (Hartl and Schneider, 2019). We found that BASP1-G2A is partially defective in tumor suppressor function which is consistent with the retention of chromatin modification and transcriptional regulatory function.

The BASP1 G2A mutation prevents-terminal myristoylation of BASP1 and abolishes its interaction with lipids. In that respect, the findings presented here raise the question of whether the lipid-binding function of wtBASP1 can be modulated to provide a regulatory mechanism for the transcriptional repression function of BASP1. Indeed, previous studies have demonstrated that phosphorylation of Serine-6 within BASP1 generates an unfavourable environment for the binding of lipids by BASP1 (Mosevitsky, 2005). We have previously reported that phosphorylation of BASP1 disrupts its transcriptional repressor activity (Toska et al., 2012). This raises the possibility that phosphorylation of BASP1 might regulate its ability to interact with different chromatin remodelling activities in the regulation of the chromatin landscape.

Our RNAseq data suggest that BASP1 can regulate the activity of several transcription factors. These include transcription factors previously identified as partners of BASP1, MYC (Hartl et al., 2009), YY1 (Santiago et al., 2021) and CTCF (Essafi et al., 2011). Several target genes of MYC and the associated MAZ protein were regulated by BASP1 which had both coactivator and corepressor function that was not dependent on the myristoylation of BASP1. This reduced dependence on BASP1 myristoylation might be due to the proposed alternative mechanism whereby BASP1 interferes with the MYC-Calmodulin interaction. In contrast, YY1 and CTCF target genes showed BASP1-dependent repression that was substantially dependent on BASP1 myristoylation. RELA and TFAP2A/C target genes were also repressed by BASP1 that was dependent on BASP1 myristoylation. Taken together, these results suggest that BASP1 modulates the transcriptional activity of several transcription factors through both lipidation-dependent and lipidation-independent mechanisms.

## Acknowledgements

This work was funded by the BBSRC so SGER (BB/T001925/1) and NIH to KFM and SGER (1R01GM098609). AEL was supported by a Wellcome Trust PhD Studentship for the Dynamic Cell Biology program (083474). Would like to thank Mutia Muna for help with the BASP1 mouse experiments.

## Author contributions

The study was conceptualized and designed by AJM, AEL, KFM and SGER. All authors performed experiments and analysed the data. AJM and SGER wrote the manuscript. All authors discussed results from the experiments and commented on the manuscript. SGER supervised the project.

## Methods

### Cell lines

K562 cells were maintained in RPMI 1640 (ThermoFisher) supplemented with 10% (v/v) fetal calf serum (Sigma-Aldrich), 1% (v/v) Pen-Strep (Sigma-Aldrich) and 1% (v/v) L-glutamine (Sigma-Aldrich). Stably transfected K562 cell line derivatives were supplemented with 1mg/ml G-418 (Sigma-Aldrich). Cells were incubated at 37°C in humidified 95% air and 5% CO2. Lipid-free media was prepared as above but using charcoal-stripped fetal calf serum (Thermofisher).

### Chromatin Immunoprecipitation

K562 cells were harvested and resuspended to a density of 1x10^6^ cells/ml in PBS. Crosslinking was initiated by the addition of formaldehyde to a final concentration of 1.42% (v/v) and incubation at room temperature for 15 mins. Crosslinking was terminated by the addition of glycine to a final concentration of 125mM glycine and incubation at 4oC for 5 mins. The cells were then harvested by centrifugation at 2000 x g for 5 mins at 4°C and then washed with PBS. Cells were resuspended and lysed in 1ml IP buffer (150mM NaCl, 50mM Tris-HCl (pH 7.5), 5mM EDTA, 0.5% (v/v) NP-40 and 1% (v/v) Triton X-100) with protease inhibitor cocktail for 15 mins on ice. After centrifugation at 2000 x g for 5 mins at 4°C, the pellet resuspended in 1ml IP buffer. The chromatin was sheared by sonication using a QSonica Q500 at 60% amplitude and the lysate was cleared by centrifugation at 12000 x g for 10 mins at 4°C. The samples were precleared by incubation for 1 hour at 4°C on a rotator with 10μl Protein G magnetic beads (ThermoFisher). The indicated antibody was incubated in 600μl IP buffer containing 1μl 10mg/ml acetylated BSA and 5μl Protein G magnetic beads for 4 hours with rotation at 4°C. 200μl of the pre-cleared chromatin was added and incubation continued at 4°C overnight. A sample of the input chromatin was stored for later decrosslinking and processing.

After incubation, the samples were collected on a magnetic rack and beads sequentially washed once in IP buffer, high salt IP buffer (500mM NaCl, 50mM Tris-HCl (pH 8.0), 5mM EDTA, 0.5% (v/v) NP-40, 1% (v/v) Triton X 100), LiCl buffer (10mM Tris-HCl (pH 8.0), 250mM LiCl, 1mM EDTA, 1% (v/v) NP-40, 1% (w/v) Sodium Deoxycholate), and TE buffer (10mM Tris-HCl (pH 8.0), 1mM EDTA). The beads were then resuspended in 100μl of PK buffer (125mM Tris-HCl (pH 8.0), 10mM EDTA, 150mM NaCl, 1% (w/v) SDS) and incubated at 65°C overnight. 1μl of 20mg/ml Proteinase K fwas then added and the samples incubated for at 55°C for 4 hours. The immunoprecipitated DNA was finally purified using the Qiaquick PCR purification kit (Qiagen).

### Quantitative PCR analysis

Total RNA was prepared using the RNeasy kit (Qiagen) and then cDNA prepared using the Iscript cDNA synthesis kit (Bio-Rad). Quantitative PCR was performed in triplicate using the iQ SYBR green master mix (Bio-Rad) on a Bio-Rad Miniopticon system. ChIP analysis was performed using promoter-specific primers as indicated and fold-enrichment calculated against a control genomic region. Melt curve analysis was performed at the end of each run. Data were collected using the BioRad-CFX Manager software. The relative expression of each target gene is expressed relative to control GAPDH. Data are expressed as mean with standard deviation (SDM). Data distribution and significance between different groups was analyzed in Excel or OriginPro 7.5 using Student’s t test.

### Immunofluorescence

K562 cell line derivatives were collected by centrifugation at 1400 x g for 3 mins and nuclei were isolated using the Nuclei EZ Prep nuclei isolation kit (Sigma-Aldrich). The nuclei were fixed in 4% (v/v) paraformaldehyde at room temperature for 15 mins, then incubated with 50mM NH4Cl for 15 mins with rotation. The fixed nuclei were then washed three times in PBS and then incubated in blocking buffer (2% (w/v) BSA, 0.25% (w/v) Gelatin, 0.2% (w/v) Glycine, 0.2% (v/v) Triton X-100 in PBS) for 1 hour with rotation at room temperature. Primary antibody was added to each sample of nuclei in PBS containing 1% (w/v) BSA, 0.25% (w/v) gelatin and 0.2% (v/v) Triton X-100 and then incubated for 1 hour with rotation at room temperature. The nuclei were then washed three times with washing buffer (0.2% (w/v) Gelatin in PBS). Fluorescent secondary antibody in PBS containing 1% (w/v) BSA, 0.25% (w/v) gelatin and 0.2% (v/v) Triton X-100 was then incubated with the nuclei for 45 mins in with rotation in the dark. After washing three times the samples were incubated with DAP solution for 10 minutes and then resuspended in a minimum volume of DABCO mounting media (Sigma-Aldrich) an applied to poly-lysine coated slides. Nuclei were viewed using a Leica SP5-II AOBS confocal laser scanning microscope attached to a Leica DM I6000 inverted epifluorescence microscope with oil 63x lens. Images were processed using ImageJ or Volocity 6.3 software.

### Click Chemistry

K-562 cells were incubated in lipid free media containing 10μg/ml myristic acid-alkyne (Avanti) for 16 hrs. Cells were then harvested and washed once in PBS and the nuclei isolated using the EZ Prep nuclei kit. The purified nuclei were then incubated with 50μM Azide-PEG3-biotin conjugate (Sigma-Aldrich) in Buffer A and added to samples. The click reaction was initiated via addition of 2mM CuBF4 in 2% (v/v) acetonitrile, and the reaction was left to proceed at 43°C for 30 mins with rotation. The nuclei were then washed in 0.1M HEPES/KOH (pH 7.4) and immediately used for either preparation of nuclear extract for immunoprecipitation, chromatin immunoprecipitation or immunofluorescence.

### Immunoprecipitation

Following completion of the click chemistry reaction, nuclear extracts were prepared by resuspending samples in one PCV of NE2 buffer (20mM HEPES pH 8.0, 1.5mM MgCl2, 25% (v/v) Glycerol, 420mM NaCl, 0.2mM EDTA, 1mM DTT and 0.5mM PMSF) and incubating on ice with regular stirring for 30 mins. Nuclear debris was pelleted by 5 mins in a microfuge at full speed. The cleared nuclear extracts were then subjected to immunoprecipitation using streptavidin magnetic beads (Invitrogen) and IP buffer (20mM HEPES pH 8.0, 100mM KCl, 0.2mM EDTA, 20% (v/v) Glycerol, 0.5mM DTT and 0.05% (v/v) NP-40) overnight at 4°C. Magnetic beads were collected and washed 3 times in IP buffer.

### Colony Formation Assay

The K562 cell line derivatives were seeded onto agar bases (0.7% (w/v) LMP Agarose, 10% (v/v) FBS, 1% (v/v) Pen Strep, 1% (v/v) L-glutamine in DMEM media supplemented with 0.15% (v/v) NaHCO3, 200μM sodium pyruvate and 800μM NaOH at a density of 2x105 cells per ml of agar. Agar dishes were fed with additional agar (2ml) on days 7 and 14 post seeding. The colony formation efficiency (%CFE) was calculated 7, 14 and 21 days post seeding via counting colonies consisting of greater than 50 cells, in a minimum of ten 15.89mm^2^ fields of view per 60mm^2^ dish. Average colony area was measured using ImageJ software. All experiments were performed in triplicate.

### Animals

The floxed BASP1 mice in a C57BL/6 background have been described before (Gao et al., 2019). Animals were cared for in compliance with the University at Buffalo Animal Care and Use Committee. BASP1fl+/− mice were mated with Krt8- Cre/ERT2+ mice (The Jackson Laboratory) to obtain BASP1fl+/−Krt8-Cre/ ERT2+ mice and then bred to obtain BASP1fl+/+Krt8-Cre/ ERT2+ mice. BASP1fl−/−; Krt8- Cre/ERT2+ or BASP1fl+/+;Krt8-Cre/ERT2− littermates were used as wild-type controls. Adult mice were gavaged with 100mg/Kg body weight of Tamoxifen dissolved in corn oil/ethanol (9:1 by volume) daily for 8 days. 7 days later the mice were euthanized and taste receptor cells harvested from the circumvallate papillae as described before (Dutta Banik et al. 2018). The mice were euthanized by carbon dioxide followed by cervical dislocation and the tongues removed. A solution containing 0.7 mg/ml Collagenase B (Roche), 3 mg/ml Dispase II (Roche), and 1 mg/ml trypsin inhibitor (Sigma-Aldrich) was then injected beneath the lingual epithelium. The tongues were then incubated in in oxygenated Tyrode’s solution for 15 minutes and then the epithelium peeled from the muscle and placed in Ca2+-free Tyrode’s. The cells were used immediately for either RNA preparation or ChIP.

### Sample preparation and sequencing

All samples were prepared in biological triplicates for V-K562, B-K562 and G- K562 cells harvested at 0.7-0.8x10^6^ cells /mL (Loats et al., 2021; Toska et al., 2012). Cells taken for ATAC-seq were treated with DNase and incubated for 30mins buffered with 25mM MgCl2 and 5mM CaCl2 prior to harvesting. Samples of 2 x10^6^ for RNA-seq and of 1 x10^5^ cells for ATAC-seq were then pelleted and snap-frozen in liquid nitrogen until library preparation. Total RNA was isolated from the cells using the RNeasy Mini Kit (Qiagen, cat 74104), RNA Integrity Number (RIN) was determined for each sample using Tape Station 4150 System (Agilent). Libraries were prepared according to manufacturer’s instructions for TruSeq Stranded mRNA Library kit (Illumina, cat #20020594) and ATAC-Seq Kit (Active Motif, cat #53150). Libraries were sequenced on Illumina NextSeq 500 in 75-nt (RNA-seq) and 42-nt (ATAC-seq) experiments in paired-end mode.

### RNA-seq and ATAC-seq data analysis

Reads were mapped to GRCh38 human reference genome using STAR(Dobin et al., 2013) (RNA-seq) and BWA-MEM(Li and Durbin, 2009) (ATAC-seq), duplicate reads were removed, uniquely mapping reads with ≤ 2 mismatches were kept for downstream analysis. RNA-seq paired reads were assigned to RefSeq transcripts using Subread and annotated with Entrez gene identifiers. ATAC peaks were called using MACS2, merged regions were defined by overlapping intervals between all samples, peak fragment densities for merged regions were used for comparative analysis and annotated with the closest gene within a 10kb window. DESeq2 was used for differential expression analysis to determine p-adjusted values (padj) and shrunken Log2 fold-change (FC) for RNA-seq genes and ATAC-seq merged regions.(Gaspar, 2018; Liao et al., 2013; Love et al., 2014)

The 5000 most differentially expressed genes for BK vs VK at padj ≤ 0.05 were analysed using iPathway to determine significantly differentially regulated networks. The ‘Impact Analysis’ was applied to underlying pathway topologies comprised of genes and their directional interactions. The probability of observing the number of DE genes in a given pathway that is ≥ to that observed by random chance (pORA) and perturbation accumulation, based on the combined perturbation of all genes within the pathway as a function of normalised change in expression and weighting in relation to gene type, (pAcc), are the principal measures of pathway enrichment. The prediction of upstream regulators is based on the enrichment of differentially expressed genes within networks of regulatory interactions, briefly, each upstream regulator *u*, the number of consistent DE genes downstream of *u*, DTI(u) is compared to the number of measured target genes expected to be both consistent and DE by chance. The z-score *Pz* is the one-tailed area under the probability density function for a normal distribution, N(0,1), *Pn* is based on the number of DE targets consistent with inhibition.(Draghici et al., 2007; Tarca et al., 2009)

## Resources and tools

The following resources were used in pathway enrichment analysis. Gene Ontology Consortium database (as at Oct 2020)(Ashburner et al., 2000).

TARGETSCAN (v7.2)(McGeary et al., 2019). miRBase (v22.1, 2020)(Kozomara et al., 2019). KEGG database (v96.0, 2020)(Kanehisa et al., 2012). Comparative Toxicogenomics Database (as at Jul 2020)(Davis et al., 2019) . BioGRID: Biological General Repository for Interaction Datasets (v4.0.189, 2020)(Szklarczyk et al., 2017).

Heatmaps were produced from RNA-seq scaled read counts using pheatmap (https://github.com/raivokolde/pheatmap), and from ATAC-seq bigwig signals, log2 normalised for BK and GK against VK, using deepTools2 (https://github.com/deeptools/deepTools) (Ramírez et al., 2016). Volcano plots were produced using EnhancedVolcano (https://github.com/kevinblighe/EnhancedVolcano). Additional plots were prepared with ggplot2 (https://github.com/tidyverse/ggplot2). Pavis2 (https://manticore.niehs.nih.gov/pavis2/)(Huang et al., 2013) and GREAT (http://great.stanford.edu/public/html/)(McLean et al., 2010) were used with default settings for annotations. g:Profiler (https://biit.cs.ut.ee/gprofiler/gost)(Raudvere et al., 2019) was used to explore gene set functional annotations. The TRANSFAC (https://maayanlab.cloud/Harmonizome/resource/TRANSFAC) and MotifMap (https://maayanlab.cloud/Harmonizome/dataset/MotifMap+Predicted+Transcription+Factor+Targets) databases were used for exploration of transcription factors and their targets, binding sites for CTCF were assigned as lost or gained in cancer according to Fang et al (2020). IGV was used to capture genome browser views (https://github.com/igvteam/igv) (Thorvaldsdóttir et al., 2013).

Oligonucleotides

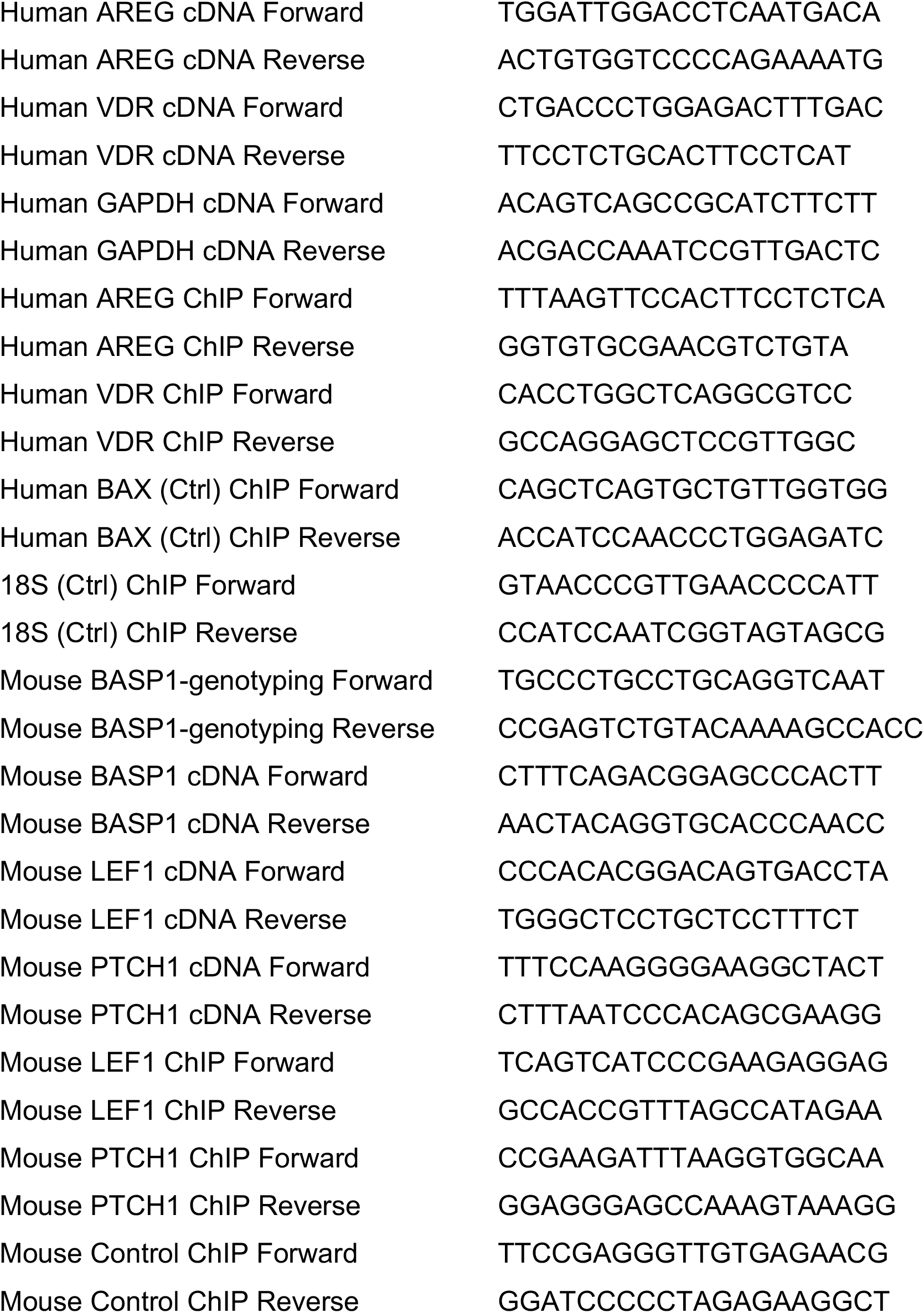

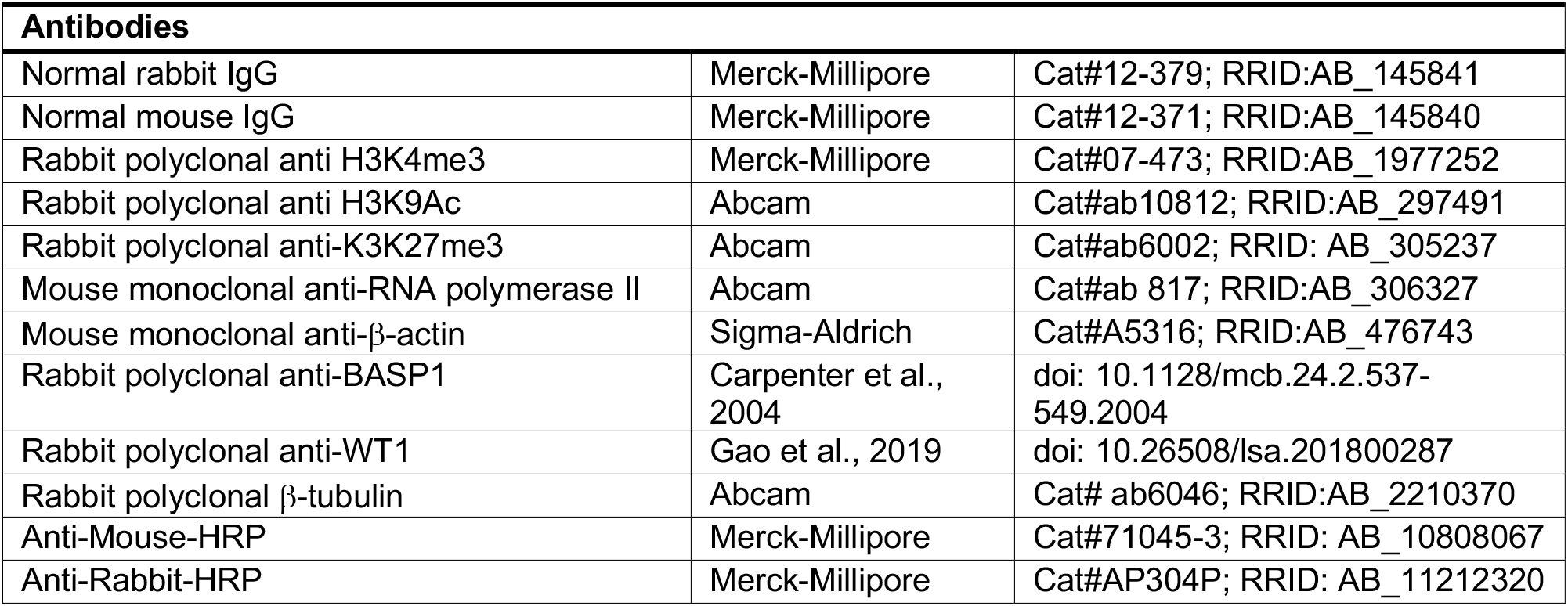

---

**Supplementary Figure 1.**
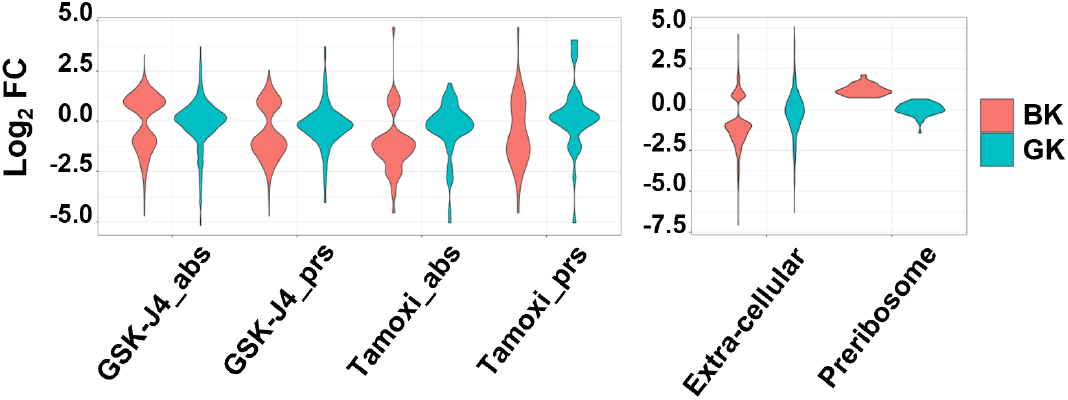
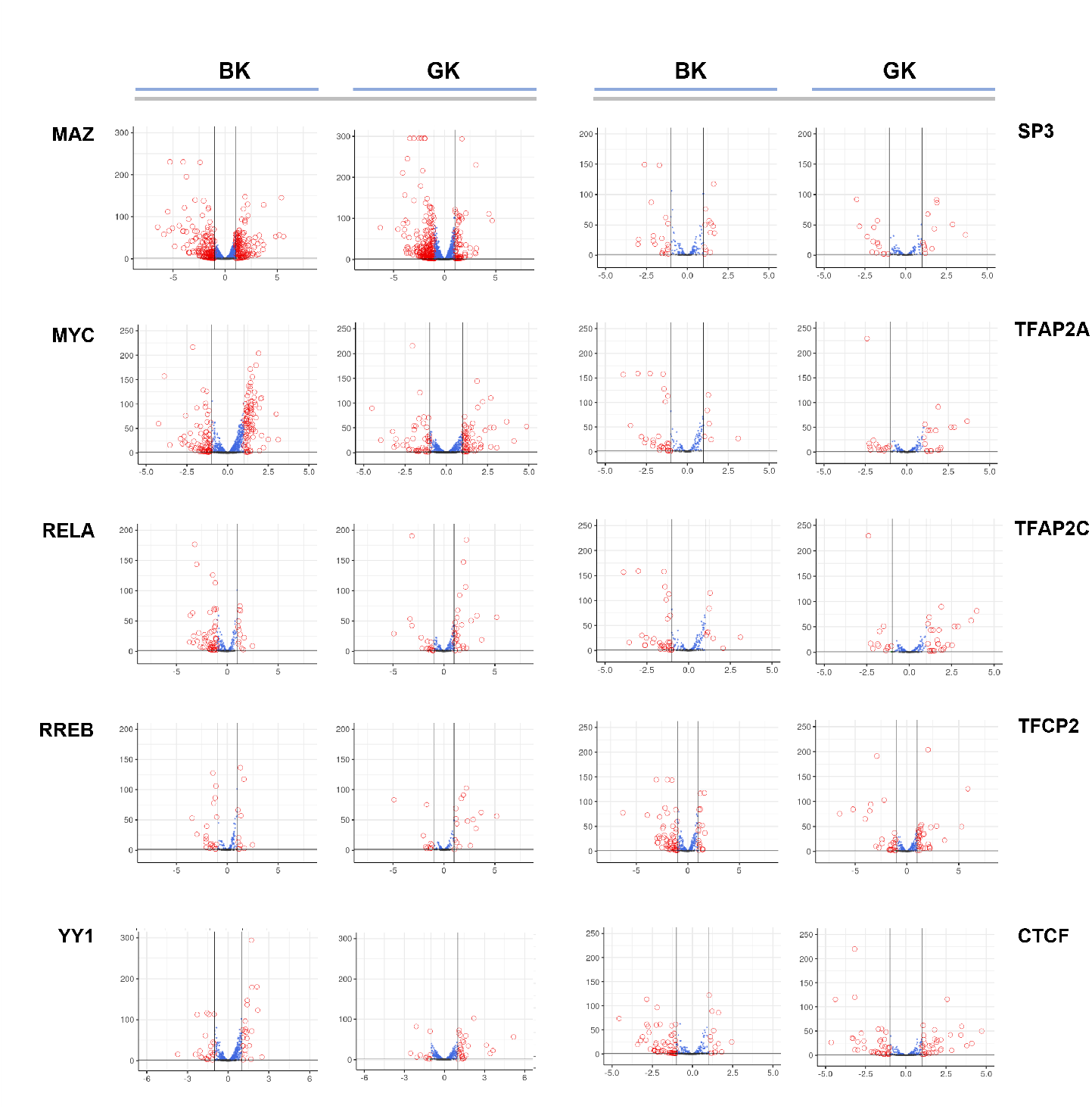
S1A: Differential gene expression for genes associated with the absence or deficiency (abs) and presence or abundance (prs) of GSK-J4 and tamoxifen (left panel), and those involved in regulation of the extracellular space and pre-ribosome processes (right panel). S1B: Functional enrichment of transcription factors (TFs) were identified from significantly differentially expressed genes (padj ≤ 0.05, FC ≥ 1, BK vs VK), YY1 and CTCF targets are also included, differential expression of TF target genes is shown in volcano plots for BK and GK.

**Supplementary Figure 2.**
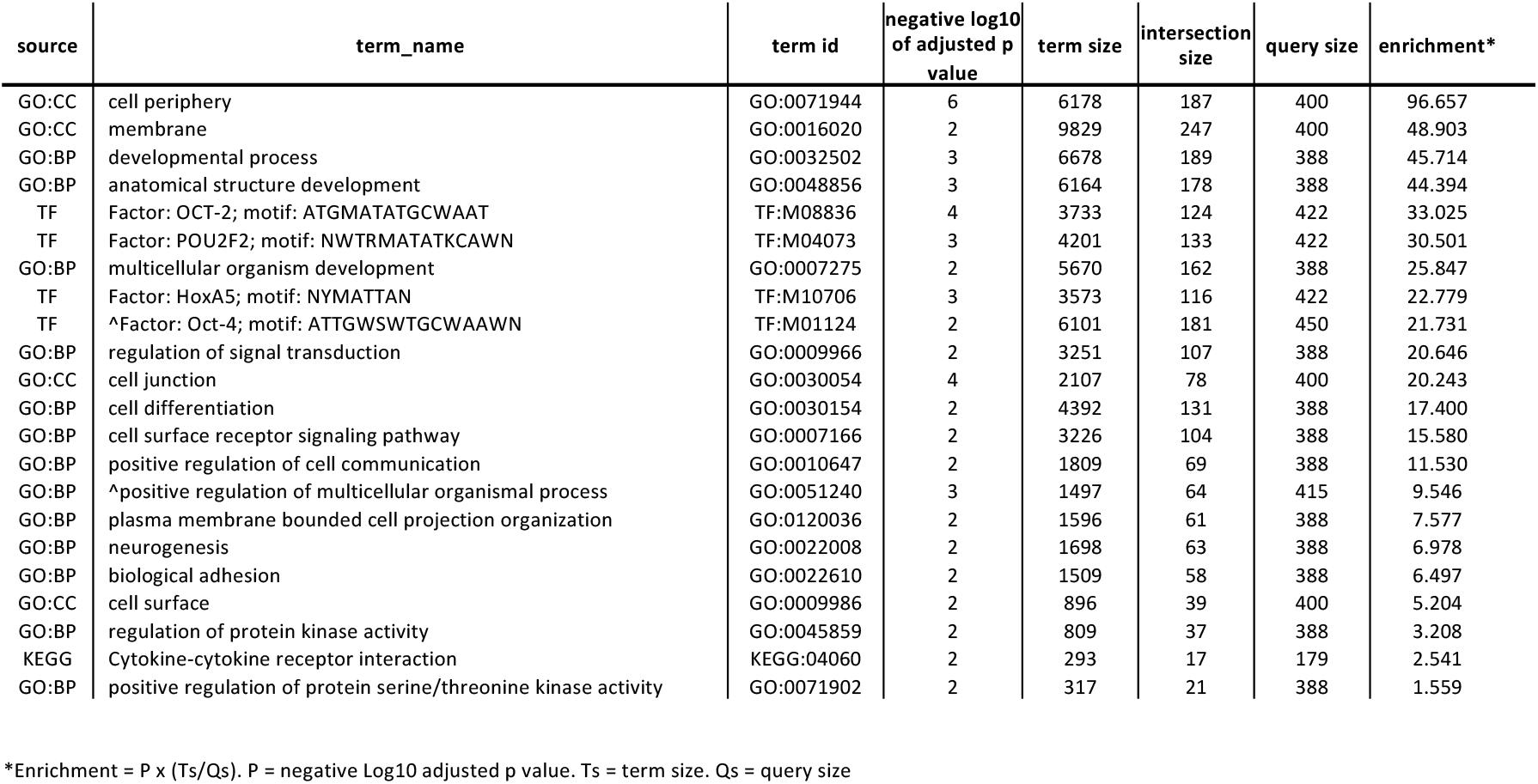
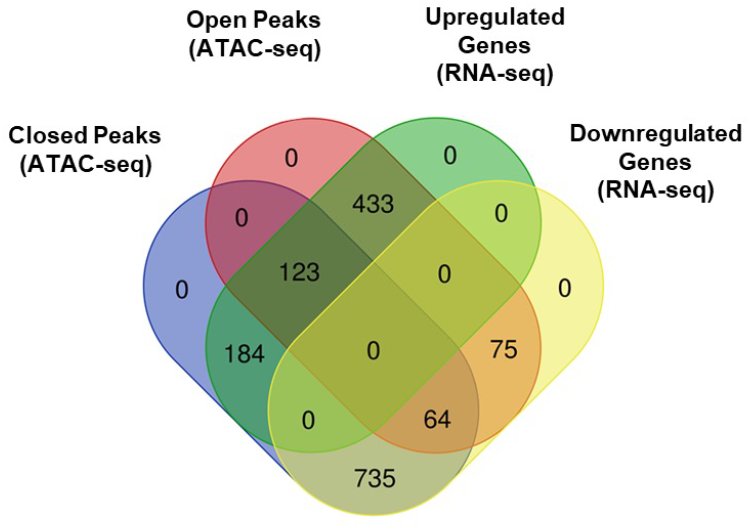
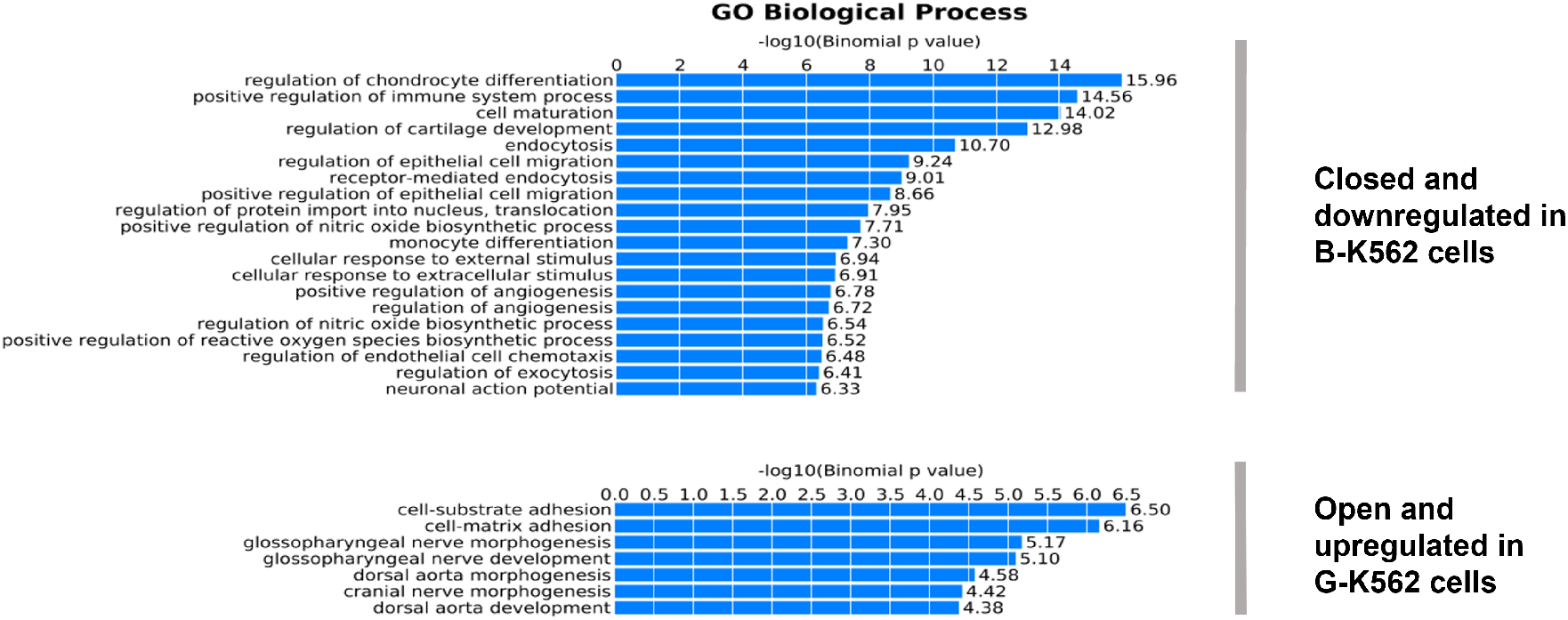
Figure S2A: Table produced from combined g:profiler results for genes (ordered and unordered analysis), and peak regions (unordered analysis only) for down regulated genes with closed chromatin (peaks and genes at padj ≤ 0.05 and FC ≥ 1, BK vs VK) (see Fig 4). Terms shown appeared in ≥ 2 of the three analyses, statistics shown are from unordered peak analysis, except where shown from unordered gene analysis ^. Data are ordered by enrichment score calculated as a function of the negative Log10 adjusted p value and term size to query size ratio. Figure S2B: Up and down regulated genes counts at open and closed chromatin regions are shown for differentially expressed peaks and genes at padj ≤ 0.05 and FC ≥ 1 for GK vs VK. See also Figure 4. Figure S2C: Functional enrichment among chromatin regions.

